# A coral peptide has bactericidal activity against a global marine pathogen, *Vibrio coralliilyticus*

**DOI:** 10.1101/2024.09.18.613810

**Authors:** Kako Aoyama, Masahiko Okai, Nobuhiro Ogawa, Riko Fukumaru, Masami Ishida, Koji Inoue, Toshiyuki Takagi

## Abstract

Scleractinian corals and their associated microorganisms, including endosymbiotic dinoflagellates and bacteria, constitute coral holobionts. Rising seawater temperatures weaken coral symbiotic relationships, thereby increasing susceptibility to infectious diseases and leading to disease outbreaks and subsequent population declines. The temperature-dependent coral pathogen, *Vibrio coralliilyticus*, poses one of the greatest threats to coral reefs because of global warming. However, coral immune defenses against this pathogen are poorly understood. We previously identified coral genes responding to *V. coralliilyticus* exposure by transcriptomic analysis of the reef-building coral, *Acropora digitifera*. Here, bioinformatic analysis identified digitiferin, a coral antimicrobial peptide (AMP), in the genome of *A. digitifera*. Recombinant digitiferin showed antibacterial activity against both Gram-positive and -negative bacteria. A 3D structural model and direct microscopic observations revealed that digitiferin damaged bacterial cell membranes by forming pores; however, initial growth inhibition tests in 1.5 % (w/v) NaCl showed no effect on pathogens. Because we found that digitiferin is secreted from epithelial cells into mucus and shows salt sensitivity, we hypothesized that it is active against pathogens in low-salt environments. We then investigated whether it showed antibacterial activity against pathogens in low-salt, moderate-osmolality conditions, using mannitol for osmoregulation instead of NaCl. The results showed that digitiferin exhibits bactericidal activity against *V. coralliilyticus* under salt-free conditions. To the best of our knowledge, this is the first study to characterize a scleractinian AMP that kills *V. coralliilyticus*. These findings contribute to a better understanding of coral immunity and may facilitate development of techniques to overcome coral *Vibrio* disease.

**Significance:** Healthy coral reefs are one of the most important marine ecosystems, supporting ∼25 % of all marine organisms. Coral reef ecosystems are threatened by coral infectious diseases, as rising seawater temperatures exacerbate pathogen infectivity. Thus, understanding coral immune systems is becoming increasingly important to protect coral reefs. This study reports the discovery of an antimicrobial peptide, digitiferin, from the reef-building coral, *Acropora digitifera*. Digitiferin is effective against Gram-positive and -negative bacteria, including the major coral pathogen, *Vibrio coralliilyticus*, and is secreted into mucus from ectodermal granular epithelial cells. As the first antimicrobial peptide known to have bactericidal activity against *V. coralliilyticus*, this discovery enhances our understanding of coral immunology and coral-pathogen interactions.

## Introduction

Coral reefs are important marine ecosystems that afford habitat for ∼25% of all marine species, although they cover <0.1% of the ocean floor. However, they are under serious threat of collapse due to large-scale bleaching, caused by rising seawater temperatures due to global climate change, and bacterial infections. Increased seawater temperatures lower the resistance of corals to diseases and enhance pathogen growth, infectivity, and virulence (1–3). Since the 1980s, the incidence of coral diseases caused by bacterial infections has increased dramatically (4, 5). *Vibrio* species are widely distributed throughout the ocean and are the major pathogenic bacteria in corals. The temperature-dependent pathogen, *Vibrio coralliilyticus*, is the best-known causative agent of bacterial bleaching, white syndrome, and tissue lysis in corals (6–8). Various virulence factors of *V. coralliilyticus* involved in motility, secretion systems, host degradation, and transcriptional regulation, including quorum sensing, are augmented by high temperatures (9, 10). However, coral immune defense mechanisms against these pathogens, including *V. coralliilyticus*, are poorly understood.

Because corals and other invertebrates lack adaptive immune systems, their innate immune responses to pathogens are important determinants of disease susceptibility (11). There are three major elements of the innate immune response: immune recognition of pathogens, intracellular signaling to activate antimicrobial responses, and effector responses (12). Whole-genome sequencing of the reef-building coral, *Acropora digitifera*, has revealed that the number of pattern recognition receptors for pathogen recognition, such as Toll-like and nucleotide oligomerization domain-like receptors, is much greater than that of *Nematostella vectensis* and *Hydra magnipapillata*, so the repertoires for innate immunity in corals are probably more complicated than those of sea anemones and hydras (13, 14). As for effectors, antimicrobial peptides (AMPs) serve important functions in the innate immune systems of both vertebrates and invertebrates. AMPs are inducible and specific for invading pathogens such as fungi, bacteria, and viruses (15). Generally, AMPs are positively charged and amphiphilic. Therefore, they are attracted to negatively charged bacterial cell membranes via electrostatic attraction and perforate bacterial cell membranes by inserting their hydrophobic moieties into lipid bilayers (16).

In corals, only two AMPs have been previously identified, damicornin from *Pocillopora damicornis* (17) and AmAMP1 from *Acropora millepora* (18), although >800 coral species have been identified worldwide. Damicornin exhibits antimicrobial activity against Gram-positive bacteria and the fungus, *Fusarium oxysporum*, but has low antimicrobial activity against Gram-negative bacteria and no activity against *Vibrio* species, including *V. coralliilyticus*. AmAMP1 shows antibacterial activity against both Gram-positive and -negative bacteria; however, it also shows very low activity against *V. coralliilyticus*. In other words, even though *V. coralliilyticus* is a well-known pathogenic bacterium of corals, immune defense systems of corals against this pathogen are completely unknown.

Here, we identified an AMP, digitiferin, among genes that respond to V. coralliilyticus using transcriptomic data of *A. digitifera* (19). To characterize digitiferin, it was heterologously expressed in *Escherichia coli*, and its antimicrobial activity against Gram-negative and -positive bacteria was demonstrated using a liquid growth inhibition assay. Immunohistochemical analysis revealed that digitiferin is secreted from granular epithelial cells onto the body surface and mucus layer of *A. digitifera*. Unexpectedly, digitiferin appeared ineffective against coral pathogenic bacteria in a growth inhibition assay in NaCl solution. Finally, we discovered that digitiferin activity varies with salinity and demonstrated that it is effective against the coral pathogen, *V. coralliilyticus*, under salt-free conditions.

## Results

### Identification of digitiferin from *A. digitifera*

To investigate novel antimicrobial peptides, short proteins (<100 AA) from genes in the coral, *A. digitifera*, responding to *V. coralliilyticus* exposure (19) that possess an N-terminal signal peptide, but lack clear homologs in the SwissProt database, were filtered based on AMP prediction models. We found one potential candidate, adig_s0027.g47, from gene models of *A. digitifera* using the genome browser of the OIST Marine Genomics Unit (https://marinegenomics.oist.jp/gallery) (13, 20). AMP prediction models suggested that the amino acid sequence of adig_s0027.g47 might possess antimicrobial activity (iAMPpred score: 0.99; CAMP score [support vector machine model]: 0.957; AntiBP2 score: 0.703), and we named it digitiferin (sequence in Fig. 1A). A MOTIF search indicated that the predicted mature peptide of digitiferin (81 amino acids behind the signal sequence) contained ShK domains, similar to the potassium channel toxin of the sea anemone (*Stichodactyla helianthus*) (21) (Fig. 1B).

**Fig. 1.**
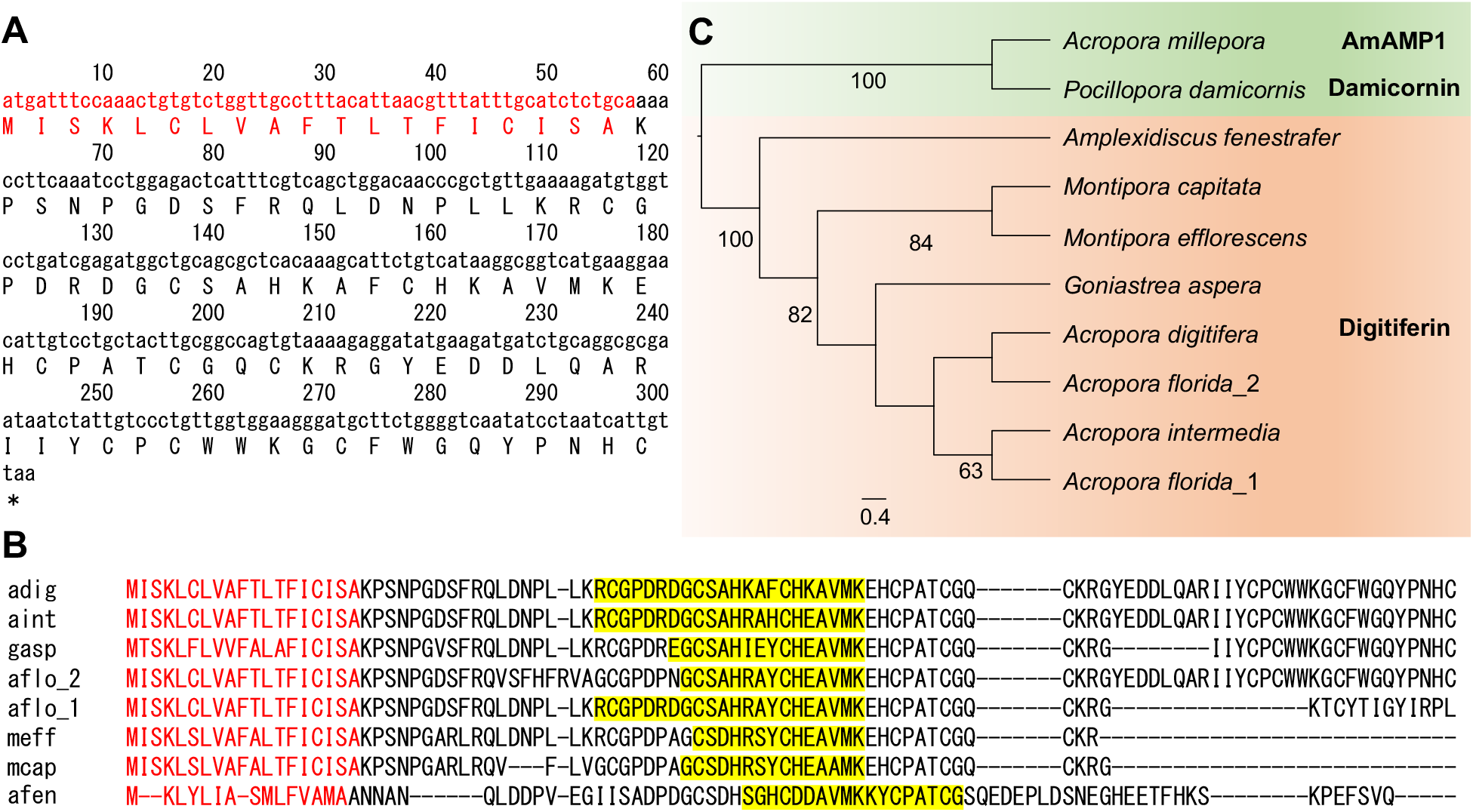
The sequence of digitiferin and those of similar genes. (A) Nucleotide and amino acid sequences of digitiferin (in red; the signal peptide predicted by SignalP5.0). (B) Alignment of digitiferin with its homologs: adi, adig_s0027.g47 from *A. digitifera*; aint, aint_s0212.g63 from *A. intermedia*; gaspa, gasp1.m3.8002.m1 from *G. aspera*; aflo_2, aflo_s0063.g40 from *A. florida*; aflo_1, aflo_s0063.g39 from *A. florida*; meff, meff_s4563.g1 from *M. efflorescensi*; mcap, OX421446.1 from *M. capitata*; and afen, evm.model.scaffold_46.36 from *Amplexidiscus fenestrafer*. Red letters indicate signal peptides. Yellow highlights indicate ShK domains predicted by a MOTIF search (https://www.genome.jp/tools/motif/). (C) Maximum likelihood phylogenetic tree showing relationships between full-length predicted digitiferin for seven corals with damicornin from *P. damicornis* and AmAMP1 from *A. millepora* as the outgroup. Bootstrap values of >60 % are indicated at nodes.

Digitiferin showed no significant homology to damicornin or AmAMP1 (based on a 1:1 alignment with NCBI BlastP, using default parameters). A tBlastN search using the genome browser of the OIST Marine Genomics Unit, Reefgenomics (http://reefgenomics.org/), and NCBI (https://www.ncbi.nlm.nih.gov/), detected genes homologous to digitiferin in several cnidarian species (*Acropora intermedia* [aint_s0212.g63], *A. florida* [aflo_s0063.g39, aflo_s0063.g40], *Montipora efflorescens* [meff_s4563.g1], *M. capitata* [NCBI sequence ID: OX421446.1], *Goniastrea aspera* [gasp1.m3.8002. m1], and *Amplexidiscus fenestrafer* [evm.model.scaffold_46.36]). The homolog in *A. florida* was tandemly duplicated with conserved synteny of neighboring genes (Fig. S1). All of these genes were aligned using MAFFT (Fig. 1B). A MOTIF search revealed that they also have ShK domains, like those of digitiferin. Fig. 1C shows results of phylogenetic analyses of precursor proteins. Although known coral AMPs, such as damicornin and AmAMP1, also have ShK domains (17, 18), maximum likelihood analysis identified seven genes similar to digitiferin as a separate group. Signal peptide sequences were highly conserved, except in *Amplexidiscus fenestrafer* (Fig. 1B). AmAMP1 has also been reported as a potential neuropeptide (18). Moreover, the conserved syntenic region of genes around digitiferin in *A. digitifera* and *A. florida* contained two genes encoding synaptotagmin, a calcium sensor that regulates exocytosis of synaptic vesicles (Fig. S1) (22).

### Heterogeneous expression and purification of digitiferin

To express mature digitiferin, cDNA fragments encoding digitiferin without the signal peptide sequence were subcloned into the pET48b(+) vector. The expression vector pET48b-digitiferin was introduced into *E. coli*, Rosetta-gami2(DE3)pLysS-competent cells. Digitiferin, with an N-terminal thioredoxin and His tag, was recombinantly expressed as a soluble fraction under isopropyl β-D-thiogalactopyranoside induction. This construct was visualized on SDS-PAGE and stained with Coomassie brilliant blue (Fig. S2). The molecular weight of recombinant digitiferin was ∼25 kDa, which is in good agreement with the molecular weight deduced from the nucleotide sequence (25,514 Da). The soluble fraction was purified using Ni-affinity chromatography and impurities were removed. By adding HRV 3C protease, recombinant digitiferin was cleaved into a thioredoxine-6×His tag (15.9 kDa) and mature digitiferin (9.6 kDa). Digitiferin was further purified using Ni-affinity chromatography to remove the thioredoxine-6×His tag. The purified solution was concentrated by ultrafiltration using an Amicon® ultra-15 centrifugal filter device with a 3-kDa cut-off to yield only digitiferin. Ten milligrams of the recombinant peptide were obtained.

### Digitiferin antimicrobial activity against *E. coli*

Antimicrobial activity of purified digitiferin was investigated preliminarily using a liquid growth inhibition assay with *E. coli* (NBRC 102203^T^). Generally, minimal inhibitory concentrations (MICs), the lowest concentrations of antimicrobial agents that completely inhibit bacterial growth in 96 well plates, as measured by spectrophotometry, and minimal bactericidal concentrations (MBCs), the lowest concentrations of antimicrobial agents required to kill 99.9 % of the final inoculum, are used to evaluate antimicrobial activity (23). Digitiferin did not exhibit antimicrobial activity against *E. coli*, according to MIC and MBC definitions; however, *E. coli* formed aggregates at the bottom of the plate in 10 µM digitiferin (Fig. S3A). Aggregates were stained with trypan blue, which stains only dead cells, and observed under a microscope (Fig. S3B and C). Edges of bacterial aggregates were stained blue, whereas their centers contained both live and dead cells (Fig. S3D), suggesting that *E. coli* formed aggregates to escape effects of digitiferin. Therefore, we established minimum aggregate-forming concentrations (MACs) to evaluate antimicrobial activity based on bacterial aggregate formation. The MAC is defined as the lowest concentration of an antimicrobial agent at which bacteria form aggregates. A liquid growth inhibition assay revealed that digitiferin has antibacterial activity against *E. coli* (MAC = 10 μM).

### Antimicrobial spectrum evaluation of digitiferin

To explore the antibacterial spectrum of digitiferin, we evaluated its activity against Gram-positive bacteria: (*Bacillus subtilis* [NBRC 13719^T^] and *Staphylococcus aureus* [NBRC 100910^T^]); Gramnegative bacteria: (*E. coli* [NBRC 102203^T^], *Pseudomonas aeruginosa* [NBRC 12689^T^], and *Serratia marcescens* [ATCC BAA-632]); and coral pathogens: (*V. coralliilyticus* P1 [LMG23696], *V. coralliilyticus* YB1 [ATCC BAA-450], *Vibrio shiloi* AK1 [ATCC BAA-91], and *S. marcescens* [ATCC BAA-632]). All coral-pathogenic bacteria used here were Gram negative and isolated from diseased corals. *Vibrio coralliilyticus* P1 was derived from corals affected with white syndrome (6), a general term for scleractinian coral disease, characterized by acute tissue lesions that often result in total colony mortality (6, 7). *Vibrio coralliilyticus* YB1 was derived from diseased *P. damicornis* and is the etiological agent of coral tissue lysis (24). *Vibrio shiloi* causes rapid and extensive bleaching of *Oculina patagonica* (25). *Serratia marcescens* is found in the intestines and feces of humans and livestock and causes white pox disease in corals (26).

A liquid growth inhibition assay showed that digitiferin exhibited antibacterial activity against Gram-positive bacteria *B. subtilis* (MBC = 10 μM, MIC = 10 μM, MAC = 5 μM) and *S. aureus* (MAC = 10 μM) and the Gram-negative bacterium *E. coli* (MAC = 10 μM) (Table 1; Fig. S4). However, it showed no antimicrobial activity against Gram-negative *P. aeruginosa* and *S. marcescens*, or coral-pathogenic bacteria (MBC > 10 μM, MIC > 10 μM, MAC > 10 μM) (Table 1; Fig. S5).

**Table 1.** MBCs, MICs, and MACs of digitiferin obtained by liquid growth inhibition assays.

| | Strain | MBC<br>(μM) | MIC<br>(μM) | MAC<br>(μM) |
| --- | --- | --- | --- | --- |
| Gram-positive<br>bacteria | *Bacillus subtilis* (NBRC13719<sup>T</sup>) | 10 | 10 | 5 |
|  | *Staphylococcus aureus* (NBRC100910<sup>T</sup>) | >10 | >10 | 10 |
| Gram-negative<br>bacteria | *Escherichia coli* (NBRC102203<sup>T</sup>) | >10 | >10 | 10 |
|  | *Pseudomonas aeruginosa* (NBRC12689<sup>T</sup>) | >10 | >10 | >10 |
|  | *Serratia marcescens* (ATCC BAA-632) | >10 | >10 | >10 |
| Coral pathogenic<br>bacteria | *Vibrio coralliilyticus* P1 (LMG23696) | >10 | >10 | >10 |
|  | *Vibrio coralliilyticus* YB1 (ATCC BAA-450) | >10 | >10 | >10 |
|  | *Vibrio shiloi* AK1 (ATCC BAA-91) | >10 | >10 | >10 |
|  | *Serratia marcescens* (ATCC BAA-632) | >10 | >10 | >10 |

MBC, minimal bactericidal concentration; MIC, minimal inhibitory concentration; MAC, minimum aggregation formation concentration.

### Predicted mechanism of action with a digitiferin structural model and observations of digitiferin-treated bacteria using SEM

To better understand the mechanism of its antimicrobial action, a structural model of digitiferin was constructed using ColabFold (Fig. 2A). Although digitiferin has an ShK-like domain, based on a MOTIF search, no structural similarity to known proteins was found using the DALI server. Digitiferin can be divided into neutral and positive regions; therefore, it was predicted that digitiferin interacts with the core and surface of bacterial cell membranes. To confirm the predicted mechanism, *B. subtilis*, which was most sensitive to digitiferin (Table 1), was used for SEM observations. *B. subtilis* was treated with 5 μM digitiferin (MAC), and SEM was performed to visualize cell morphology and membrane integrity. Controls without digitiferin treatment had smooth, intact surfaces (Fig. 2B). Conversely, digitiferin treatment caused significant damage to bacterial cells (Fig. 2C). Large dimples formed on the cell surfaces of *B. subtilis*. Furthermore, untreated *B. subtilis* were approximately 2.6 μm in diameter, while those treated with digitiferin were reduced to approximately 1.8 µm. Additionally, cytoplasmic membrane surfaces appeared rough and were covered with numerous blebs (Fig. 2D and 2E). These results indicate that digitiferin acts on bacterial cell membranes.

**Fig. 2.**
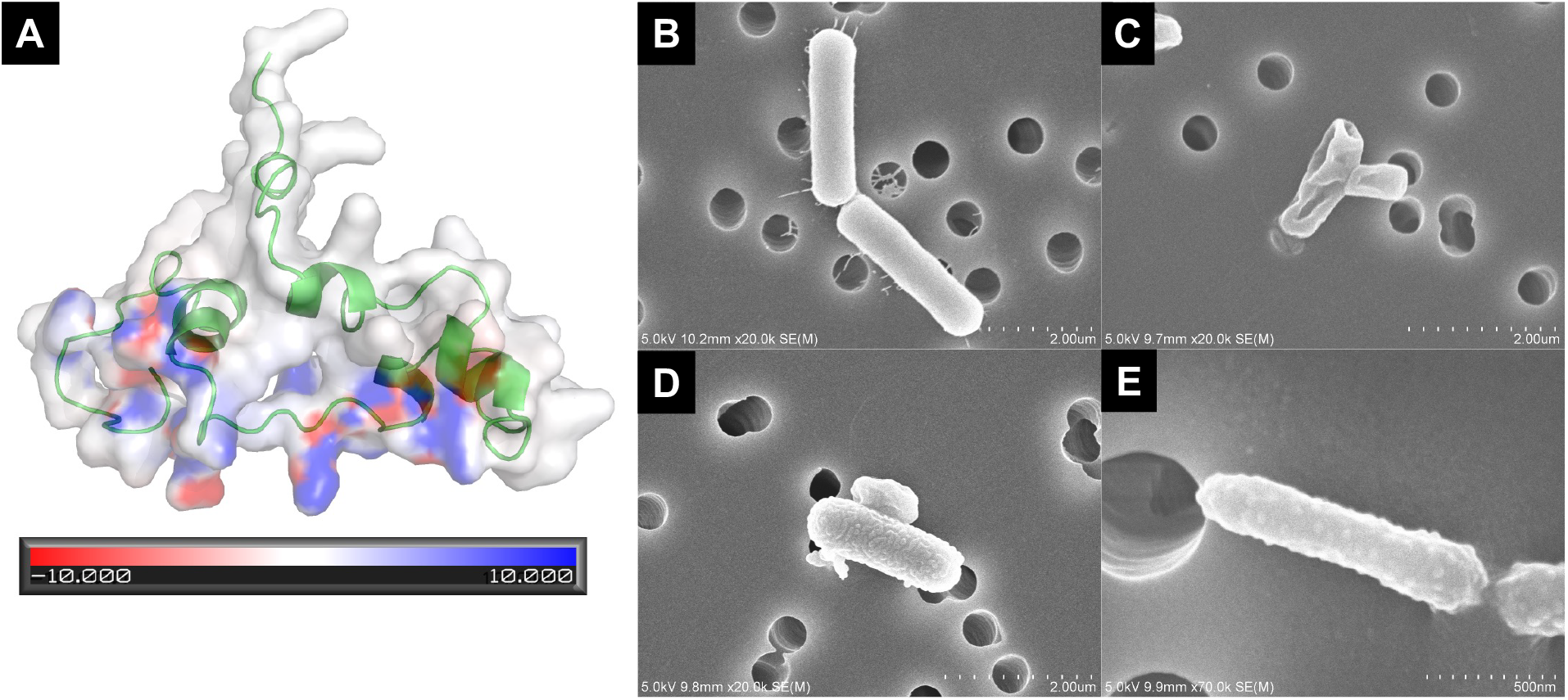
Predicted mechanism of action using a structural model of digitiferin and SEM of digitiferin-treated bacteria. (A) Structural model of digitiferin (residues 20–100). The ribbon diagram of digitiferin is colored green. The ±10 kT/e electrostatic potential is plotted on the solvent-accessible surface. White, blue, and red colored surfaces show neutral, positive, and negative potentials, respectively. Digitiferin could be divided into neutral and positive potential regions, which would interact with the cell membrane core and surface, respectively. (B) *Bacillus subtilis* with buffer. (C–E) Digitiferin-treated *B. subtilis*. (C) Large dimples formed on cell surfaces. (D) Roughness observed on cell surfaces. (E) Numerous vesicles covered the bacteria.

### Histological analysis of digitiferin expression in coral tissue

For histological characterization of cells expressing digitiferin at the protein level, a polyclonal antibody against digitiferin (anti-digitiferin antibody) was developed using 7.0 mg purified digitiferin as an antigen (Biologica, Nagoya, Japan). A dot blot analysis showed that the antibody retained distinct immunoreactivity against 10 ng of the antigen peptide, even with a 1/6,000 antibody dilution (Fig. S6), indicating that the antibody was sensitive to the antigenic peptide. Apical branches from coral colonies were mounted vertically, and coral fragments were prepared (Fig. 3A and 3B). Localization of digitiferin in coral fragments was investigated. Positive signals were detected in the coenosarc, connective tissue between polyps in a coral colony (Fig. 3C), and were particularly localized in granular epithelial cells in the ectoderm (Fig. 3D). Digitiferin is secreted by granular epithelial cells into coral mucus (Fig. 3E), but no signals were detected in endoderm. Guinea pig serum IgG from non-immunized animals was used as a negative control at the same dilution as the primary antibody. Pre-adsorption of anti-digitiferin was also performed and used as the primary antibody. All negative controls showed substantially abolished or decreased signals (Fig. 3F, Fig. S7). Taken together with the fact that digitiferin has an N-terminal signaling peptide for extracellular secretion, these results suggest that it is secreted into the mucus from granular epithelial cells in the coenosarc ectoderm.

**Fig. 3.**
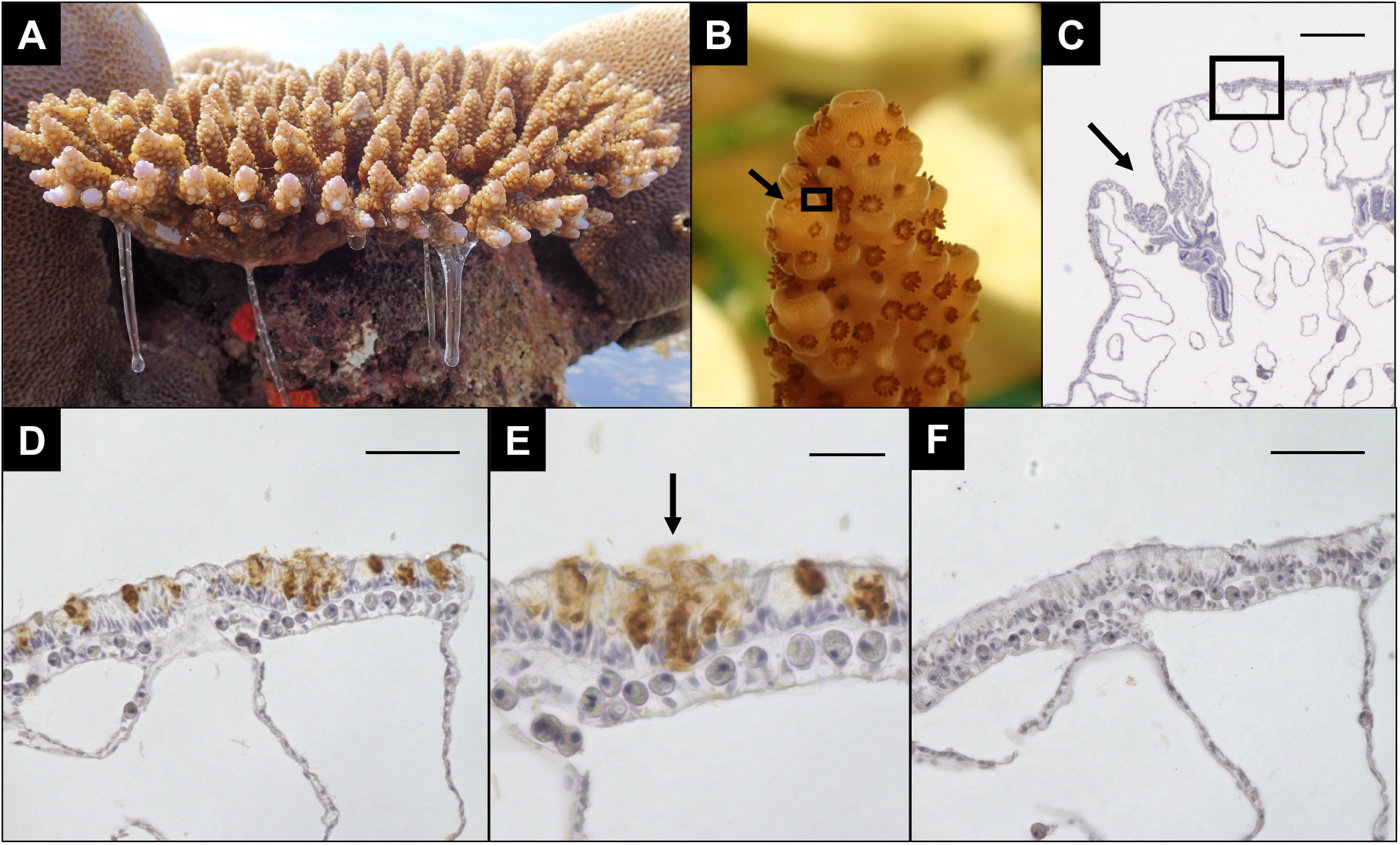
*Acropora digitifera* secreting mucus, coral fragments, and immunohistochemistry of a coral tissue section. (A) A colony of *A. digitifera* secreting mucus at low tide. (B) Coral fragments from the colony. (C) Overall view of a coral tissue section (scale bar, 200 μm). The arrow indicates a polyp. The boxed region indicates the coenosarc (B, C). (D) Immunohistochemistry with anti-digitiferin antibody (scale bar, 50 μm). (E) Higher magnification views of D. The arrow indicates extracellular secretion of digitiferin (scale bar, 20 μm). (F) Immunohistochemistry with a guinea pig serum IgG from a non-immunized animal, as the negative control (scale bar, 50 μm). Sequential sections were used for analysis and identification in D and F.

### Salt sensitivity of digitiferin

Digitiferin showed antibacterial activity against Gram-positive bacteria, *B. subtilis* and *S. aureus*, and the Gram-negative bacterium, *E. coli*, but antibacterial activity against coral pathogenic bacteria was not evident (Table 1). These results suggest that coral pathogens resist digitiferin or that other factors prevent its antimicrobial activity. Hydramacin-1 from *Hydra magnipapillata* is one of the most powerful cnidarian AMPs and is effective against Gram-positive and -negative bacteria (27); however, it loses its antimicrobial activity in 50 mM NaCl (0.29% [w/v]) (28). Furthermore, immunohistochemical analysis suggested that digitiferin is secreted into mucus; therefore, we compared the sodium ion concentrations of coral mucus and seawater in coral habitats.

Colonies of *A. digitifera* are exposed at the surface during low tide, and their secreted mucus collected by syringe aspiration (Fig. 3A). The sodium ion concentration in coral mucus was 264 mM (Fig. S8). In comparison, seawater at Sesoko Island, the sampling site for the coral *A. digitifera*, contained 449 mM. Because collected mucus was contaminated with seawater, we inferred that the NaCl concentration in the mucus immediately after secretion from the coral is even lower. Therefore, we hypothesized that digitiferin loses its antimicrobial activity in the presence of NaCl at the molarity of the culture medium of coral-pathogenic bacteria. We investigated salt sensitivity of digitiferin using a liquid growth inhibition assay in *B. subtilis*. The salinity of the medium was varied from 0 to 1.0% (w/v), and we found that digitiferin exhibited antimicrobial activity at NaCl concentrations ≦0.1% (w/v) (Fig. 4A), indicating that it is salt sensitive.

**Fig. 4.**
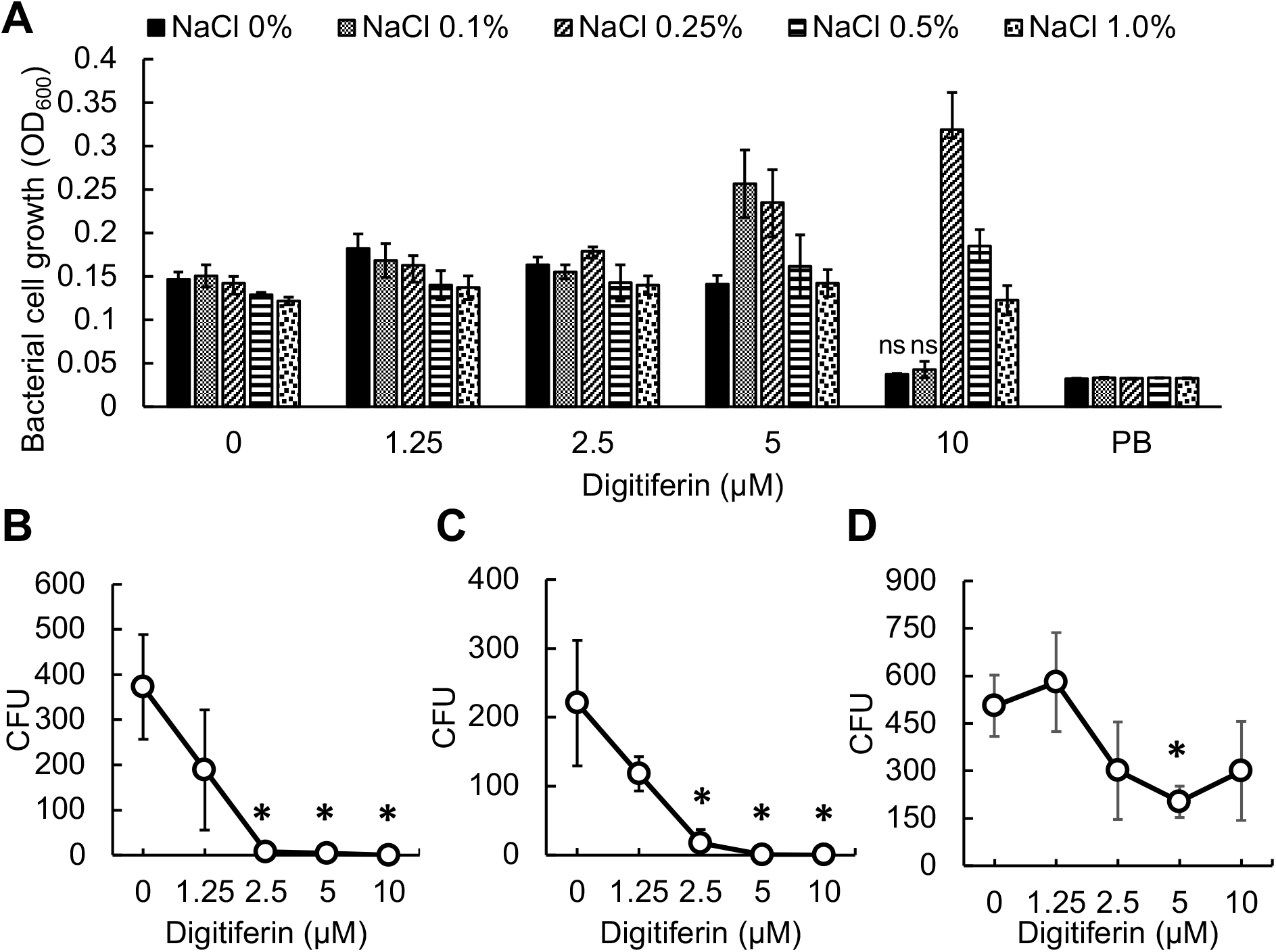
Measurements of digitiferin salt sensitivity in a liquid growth inhibition assay with *B. subtilis* and antimicrobial activity against coral-pathogenic bacteria in low-salt, moderate-osmolality conditions. (A) Growth of *B. subtilis* was monitored by measuring the OD_600_. Salinity of antimicrobial assays varied from 0 to 1.0 % (w/v) NaCl. The digitiferin concentration varied from 0 to 10 μM. PB: poor broth without *B. subtilis*. All data are presented as means ± standard error of the mean, *n* = 3. Statistical analysis between each sample and the negative control (PB) was conducted using t-test. ns shows non-significant (*P* values more than 0.05) compared to the negative control, indicating antimicrobial activity. (B–D) In the PB-mannitol medium, antimicrobial activity was determined by testing various concentrations of digitiferin. Antimicrobial activity was evaluated by plating all wells onto Marine Agar plates, and CFU counts after incubation at 30 °C for 24 h. All data are given as means ± standard error of the mean, *n* = 3. Statistical analysis between each sample and the negative control (0 μM digitiferin) was conducted using t-test. *P* values less than 0.05 were considered statistically significant. Significant differences are represented by asterisks. (B) *V. coralliilyticus* P1, (C) *V. coralliilyticus* YB1, and (D) *V. shiloi* AK1.

### Digitiferin antimicrobial activity against coral-pathogenic bacteria

We then re-evaluated digitiferin’s antimicrobial activity against coral pathogenic bacteria under salt-free conditions. As *Vibrio* spp. can survive in 1.0–7.0% NaCl (w/v) (24), we used 18.5% (w/v) mannitol, with an osmolality corresponding to 3.5% NaCl, as the exposure medium. Mannitol is a sugar alcohol abundant in marine ecosystems, and is a carbohydrate in brown algae (29). The assay was performed by exposing the bacteria to digitiferin for 3 h, the period during which *Vibrio* spp. can survive in the exposure medium. After 3 h, the bacterial solution was applied to a Marine Agar 2216 plate, and digitiferin activity was evaluated by counting the number of colonies. The negative control without digitiferin contained 200–400 CFUs (colony forming units) of *Vibrio* spp. In contrast, *V. coralliilyticus* P1 and YB1 treated with 2.5 μM digitiferin decreased to 8 and 17 CFUs, respectively, and were completely killed by 10 μM digitiferin (Fig. 4B and 4C). Digitiferin showed no antimicrobial activity against *V. shiloi* (Fig. 4D).

## Discussion

### Broad digitiferin antimicrobial spectrum, including on *V. coralliilyticus*

Here, we identified a novel AMP, digitiferin, from the reef-building coral, *A. digitifera*, among genes that respond to *V. coralliilyticus*, based on several antimicrobial peptide characteristics (Fig. 1A). In liquid growth inhibition assays, digitiferin treatment caused both Gram-positive (*B. subtilis*, and *S. aureus*) and Gram-negative (*E. coli*) bacteria to form aggregates (Table1 and Fig. S3). Such aggregations are considered ineffective by conventional AMP activity assays, such as MBC and MIC. However, aggregates of digitiferin-treated bacteria were positively stained with trypan blue, indicating bacterial death (Fig. S3B) and showing that bacterial aggregation could be an indicator of antimicrobial activity. Therefore, here, the MAC was established as the minimum concentration of AMPs required for bacteria to form aggregates. This enabled us to broadly evaluate antimicrobial activity of digitiferin.

Digitiferin exhibited the highest activity against *B. subtilis. Bacillus* spp, including *B. subtilis*, are often detected in the coral microbiome; however, their functions remain unclear (30). Colonies of *A. digitifera* are commonly observed on reef tops, and it is assumed that terrestrial bacteria are frequently transported to these areas by rivers and rainfall. Since digitiferin has antimicrobial activity against bacteria of terrestrial origin—*B. subtilis, S. aureus*, and *E. coli*—it is likely to be involved in defense against terrestrial bacteria. Although digitiferin showed antimicrobial activity against terrestrial bacteria, it was inactive against *V. coralliilyticus* P1, *V. coralliilyticus* YB1, *V. shiloi* AK1, and *S. marcescens* in 1.5% (w/v) NaCl (Table1). However, in an experiment with *B. subtilis*, we found that digitiferin lost its antimicrobial activity at concentrations >0.1% (w/v) NaCl (Fig. 4A). Therefore, we performed antimicrobial assays in a salt-free mannitol solution with an osmolality corresponding to 3.5% NaCl. Consequently, digitiferin exhibited strong antibacterial activity, killing 100% of *V. coralliilyticus* P1 and YB1 at 10 μM (Fig. 4B and C). AMPs with sufficient antimicrobial activity against pathogenic bacteria have yet to be reported, and to the best of our knowledge, this is the first report of a coral AMP with bactericidal activity against *V. coralliilyticus*, an opportunistic coral pathogen (31, 32). *Vibrio coralliilyticus* exists in healthy corals, but only invades host tissues when corals are immunodeficient or being attacked by excessive numbers of pathogenic bacteria (33). In our previous work, the digitiferin gene was upregulated by high concentrations (1.0 × 10^7^ cells/mL) of *V. coralliilyticus* (19). These results suggest that when the number of *V. coralliilyticus* in a coral exceeds a certain level, digitiferin may be expressed for defense purposes.

### Digitiferin *in vivo* expression and active sites in coral tissue

Histological analysis showed that digitiferin was secreted into the mucus from ectodermal granular epithelial cells of the coenosarc (Fig. 3C). Although damicornin from *P. damicornis* is also expressed in similar cells (17), digitiferin is the first AMP expressed in the coral coenosarc. AMP expression in granular epithelial cells has been reported in both vertebrates and invertebrates, promoting apical release of AMP in the mucus and its involvement in mucosal defense and prevention of pathogen invasion (17). Moreover, digitiferin contains an N-terminal signal peptide for extracellular secretion, and its amino acid sequence is highly conserved in corals (Fig. 1B). These findings suggest that other corals also secrete digitiferin homologs into their mucus. As it is expressed in ectodermal cells of every coenosarc between polyps of the coral colony, corals may activate an immune response throughout the whole colony against pathogens.

Scleractinian corals, including *A. digitifera*, which inhabit the upper part of reefs, secrete more mucus than other species and soft corals. They increase mucus production under stressful conditions such as exposure to air due to low tides or pathogen infection (12). Mucin is the main component of coral mucus and its physical and chemical characteristics differ completely from those of seawater. This polymeric glycoprotein provides high viscoelasticity and tensile strength, keeping the coral hydrated and trapping pathogens that attempt to invade the tissues. The sodium ion concentration in the mucus is much lower than that in seawater (Fig. S6), suggesting that the mucus just above the coral epidermis has an even lower NaCl concentration. As *Vibrio* spp. cannot survive in environments with limited salinity (24), the low salinity of mucus could be a major barrier for them. Furthermore, immune response-related genes, including digitiferin, were upregulated in response to *V. coralliilyticus* infection (19), and digitiferin showed bactericidal activity against *V. coralliilyticus* under salt-free conditions (Fig. 4B and C). These findings suggest that digitiferin is secreted under low-salt conditions, such as in mucus, and serves as a first-line defense mechanism to prevent invasion of pathogenic bacteria into coral tissues.

### The role of digitiferin in infectious disease susceptibility

Pathological agents cause disease only in certain corals, satisfying Koch’s postulates, as in the case of white pox (26), even when multiple coral species inhabit the same habitat. However, underlying mechanisms remain largely unknown. For instance, *V. coralliilyticus* (causing bacterial bleaching, tissue lysis, and white syndrome of *P. damicornis* and *Motipora aequituberculata*), *V. shiloi* (bacterial bleaching of the Mediterranean coral, *O. patagonica*), and *Aspergillus sydowii* (aspergillosis of Gorgonian corals) cause diseases in specific corals. We found digitiferin only in *A. digitifera, A. florida, A. intermedia, G. aspera, M. efflorescensi, M. capitata*, and *Amplexidiscus fenestrafer* (Fig. 1); therefore, it may be directly involved in disease susceptibility. Recently, stony coral tissue loss disease (SCTLD) has caused widespread coral loss along the Florida coral reef and devastation in various Caribbean regions (34). Over 20 species of coral are affected by SCTLD, with those from the family Meandrinidae and subfamily Faviinae the most severely damaged, while *Acropora* corals are unaffected (35). Although the etiological agent responsible for SCTLD remains unidentified, a toxic zinc-metalloprotease produced by *V. coralliilyticus* was detected in SCTLD-affected corals, suggesting that this bacterium exacerbated SCTLD lesions (36). We used *V. coralliilyticus, V. shiloi*, and *S. marcescens* to evaluate the antimicrobial spectrum of digitiferin and found that it shows antibacterial activity against *V. coralliilyticus*. Since infectious diseases involving *V. coralliilyticus* have not been reported in corals possessing digitiferin homologs, these results suggest that these corals protect themselves from *V. coralliilyticus* infections through the antimicrobial activity of digitiferin. Collectively, these findings show that immune effector genes that are present in only some species, e.g. digitiferin, may determine resistance and susceptibility to disease in corals.

### Digitiferin and its relationship to toxins

tBlastN searches of various databases revealed genes similar to digitiferin in several cnidarians. Interestingly, these genes had highly conserved sequences, including conserved signal peptides among species of *Acropora* and *Montipora* (Fig. 1B). This suggests that even minor sequence changes may affect its function, although it is possible that horizontal transmission has occurred since speciation. Motif searches revealed that coral AMPs possess a common ShK domain (Fig. 1B), also found in the potassium channel toxin of the sea anemone, *S. helianthus* (21). However, the amino acid sequence of digitiferin is distinct from that of known coral AMPs such as AmAMP1 and damicornin (Fig. 1C). Cnidarians, including corals, are equipped with unique venom organs known as nematocysts. Coral mucus not only serves as the first line of defense, but also traps particles in the water for predation (37). Digitiferin, secreted with mucus, may also be effective against planktonic predation. Several Cnidarian ShK domain-containing proteins, such as neurotoxin classicolin-I, exhibit both toxicity and AMP activity (38). The fact that drosomycin, an antifungal AMP in *Drosophila*, displays toxic activity suggests that toxins and AMPs share a common evolutionary origin in many organisms (39, 40). *Nematostella* has two paralogous ShK-like genes, ShK-like 1 (a component of nematocyst venom) and its paralog ShK-like 2 (a neuropeptide localized in neurons) (41). Intriguingly, we found that the gene of *A. florida* was duplicated (aflo_s0063.g39 and aflo_s0063.g40) and that the duplicates may have separate functions as neuropeptides and toxins, such as ShK-like1 and ShK-like2 in *Nematostella* or increase expression levels of digitiferin to create a more robust defense system.

### Harmonic regulation of pathogenic bacteria in coral mucus

In hydra, a cnidarian model organism, a secreted neuropeptide, NDA-1, with antimicrobial activity, controls the microbiome on the body surface (42). NDA-1 was detected in protrusions of the ectodermal surface, suggesting that the peptide was secreted into the mucus. In corals, mucus is a nutrient source for diverse microorganisms, and its density is higher than that of the surrounding seawater (43); however, its control mechanisms are largely unexplored. Digitiferin may regulate the mucus microbiome. In addition to AMP, regulation of bacterial quorum sensing, a means of cell-to-cell communication between bacteria dependent on population density, is another important strategy for controlling the microbiome. *Vibrio* spp. isolated from coral mucus produce the self-induced signaling molecule acyl homoserine lactone (AHL), which is involved in quorum sensing (44). AHL-mediated quorum sensing regulates expression of bacterial genes involved in various functions, including toxin expression, biofilm formation, and escape from host immune responses. Therefore, it has become apparent that AHL molecules are closely related to the survival and virulence of most bacteria (45). More recently, the quorum-quenching hydrolase AmNtNH1, which inactivates AHL, was found in the coral, *A. millepora*, and was induced by immune challenges in adult corals (46). These insights suggest that AMPs and quorum-quenching hydrolases work in concert in the mucus. That is, pathogenic bacteria weakened by quorum quenching may be eliminated in the mucus by AMPs, including digitiferin. However, further studies are needed to clarify how AMPs and quorum-quenching hydrolases interact cooperatively with pathogenic and symbiotic bacteria to regulate the coral microbiome.

## Conclusions

Rising seawater temperatures caused by global warming lower the resistance of corals to diseases and increase pathogen virulence, thereby increasing the frequency of disease outbreaks. Pathological agents cause disease only in certain corals, satisfying Koch’s postulates, even when multiple coral species/genera inhabit the same coastal region; however, the underlying mechanisms remain largely unknown. To manage disease outbreak risks in each coral species, underlying mechanisms urgently need to be identified. Here, we provided mechanistic insight into interactions between corals and the global marine pathogen, *V. coralliilyticus*, and explained mechanisms likely affecting disease susceptibility. To the best of our knowledge, this is the first study to characterize a scleractinian AMP that kills *V. coralliilyticus*, an incremental advance that enhances our understanding of coral immunology. AMPs are not only potential therapeutic agents for infectious diseases, but genes encoding them can also be applied as marker genes for diagnosing disease states; therefore, future studies of coral immunity can further contribute to the prevention and management of disease outbreaks.

## Materials and Methods

### Identification of the gene encoding digitiferin

To identify novel antimicrobial peptides, we searched for short proteins (<100 AA) among genes of the coral *A. digitifera* that are upregulated in response to *V. coralliilyticus* exposure (19). Specifically, we sought peptides that possessed N-terminal signal peptides predicted by SignalP 5.0 (https://services.healthtech.dtu.dk/services/SignalP-5.0/), but that lacked obvious homologs in the SwissProt database. These were further filtered based on the following AMP prediction models: iAMPpred (http://cabgrid.res.in:8080/amppred/), CAMP (http://www.camp.bicnirrh.res.in/prediction.php),and AntiBP2 (http://crdd.osdd.net/raghava/antibp2/help.html). We used gene models of *A. digitifera* (version 2.0) from the genome browser of the OIST Marine Genomics Unit (https://marinegenomics.oist.jp/gallery).

### Other materials and methods are provided in the Supplementary information

## Acknowledgements

This work was supported by JST ACT-X (Grant Number JPMJAX20B9), JSPS KAKENHI (Grant Numbers 21K05768, 23KJ0602, 24K08657, and 24K01848), and JST SPRING (Grant Number JPMJSP2108) in Japan. We thank Dr. Shikina Shinya and Dr. Chiu Yi-Ling (Institute of Marine Environment and Ecology, National Taiwan Ocean University) for technical advice with histological analysis.

