## Supporting_Information for "A coral peptide has bactericidal activity against a global marine pathogen, *Vibrio coralliilyticus*"


**This PDF file includes:**

Supporting text
Figures S1 to S8
Table S1
SI References

**Materials and Methods**

**Total RNA extraction from coral and cDNA cloning**

Total RNA was extracted from planula larvae, as previously described (1), to avoid contamination by endosymbiotic microalgae. cDNA was synthesized using the Superscript III First-Strand Synthesis System for RT–PCR (Thermo Fisher Scientific, Massachusetts, USA) with oligo dT primers using 500 ng of total RNA. KOD Plus Neo DNA polymerase (TOYOBO, Osaka, Japan) was used for PCR, and reaction mixtures were prepared according to the manufacturer's protocol. Primers used for PCR are listed in Table S1. The 10 × A-attachment mix (TOYOBO) allowed amplified products to acquire dA overhangs at 3'-ends. Products with 3'-dA overhangs were cloned directly into the T-vector pMD19 (Takara Bio, Kusatsu, Japan) using a Mighty Mix DNA Ligation Kit (Takara Bio). Digitiferin cDNA was amplified using M13 primers. Sequences of these amplicons were determined using an ABI PRISM 3130xl Genetic Analyzer (Applied Biosystems, Waltham, MA, USA). Nucleic acid sequences were analyzed for alignment with the *A. digitifera* genome (2, 3).

**Phylogenetic analysis**

Candidate digitiferin homologs were identified using tBlastN. The predicted *Acropora digitifera* precursor protein was used as a query against the following databases: *G. aspera* and *Amplexidiscus fenestrafer* from Reefgenomics (http://reefgenomics.org/, accessed March 17, 2023), *A. digitifera*, *A. intermedia*, *A. florida,* and *M. efflorescens* from the OIST Marine Genomics Unit web site (https://marinegenomics.oist.jp/gallery, accessed March 17, 2023), and *M. capitata* From NCBI (https://www.ncbi.nlm.nih.gov/, accessed March 17, 2023). tBlastN homology searches were conducted with a cut off of $1e^{-5}$. Protein domains of digitiferin and its homologs were searched using MOTIF (https://www.genome.jp/tools/motif/). A maximum likelihood molecular phylogenetic tree was constructed using amino acid sequences of all seven putative homologs, damicornin, and AmAMP1. Amino acid sequences of full-length proteins were aligned using MAFFT v7.450, and gaps were removed using trimAl v1.2. For tree construction RAxML v7.2.6. with the PROTCATWAG model was used (4). Bootstrap values were calculated based on 1000 replicates.

**Construction of the expression plasmid**

To understand the antimicrobial activity of digitiferin, in these experiments, we used the pET48b system in *E. coli* cells. cDNA encoding digitiferin was isolated from planula larvae. To express mature digitiferin, the cDNA fragment encoding digitiferin without the signal peptide sequence was inserted into the pET48b plasmid. Procedures were as follows: ORFs of digitiferin genes were amplified by PCR using KOD Plus Neo DNA polymerase (TOYOBO). Primers used are listed in Supplementary Table S1. PCR products were double-digested with *EcoR* I and *Sal* I and subcloned into the pET-48b vector. *E. coli* DH5α with recombinant plasmids was transformed into Luria-Bertani (LB) medium supplement with 30 mg/L kanamycin and grown at 37 ℃ till mid-logarithmic phase. A QIAprep Spin Miniprep Kit (QIAGEN, Hilden, Germany) was used to extract plasmids according to the manufacturer's protocol. Correctness of recombination was verified using DNA sequencing (GENEWIZ, Inc. Tokyo, Japan).

**Expression and purification of recombinant digitiferin**

The protein expression plasmid was transformed into *E. coli* Rosetta-gami2 (DE3) pLysS-competent cells (Merck Millipore, Massachusetts, USA). When the $OD_{600}$ reached 0.8, isopropyl β-D-thiogalactopyranoside (IPTG) was added to the culture at a final concentration of 500 μM to induce protein expression. Cell growth continued at 20℃ for 24 h. Cells were harvested by centrifugation and resuspended in sonication buffer (20 mM Tris-HCl, pH 8.0, 600 mM NaCl, and 5 mM imidazole). The cell suspension was then sonicated and centrifuged to isolate the supernatant containing the target protein. It was subjected to affinity purification using $Ni^{2+}$-nitrilotriacetate (Ni–NTA)–agarose resin (Cytiva, Tokyo, Japan). The column was equilibrated with 25 mL buffer (20 mM Tris-HCl, pH 8.0; 600 mM NaCl; 5 mM imidazole) and washed with washing buffer (20 mM Tris-HCl, pH 8.0; 600 mM NaCl; 20 mM imidazole). Elution was performed using an elution buffer (20 mM Tris-HCl, pH 8.0, 600 mM NaCl, and 200 mM imidazole). Eluted fractions were dialyzed with 1 L buffer (20 mM HEPES, pH 7.0, 600 mM NaCl) using a Spectra/Por 3 Membrane (MWCO 3.5 kDa, Spectrum Laboratories, Inc., Massachusetts, USA) at 4 ℃ for 16 h. After addition of HRV 3C protease, digitiferin was completely released from the fusion protein at 4 ℃ for 48 h. To separate the tag and digitiferin, affinity purification was performed again. Eluted fractions were concentrated by ultrafiltration using an Amicon Ultra-15 Centrifugal Filter Device (Merck Millipore) with a 3 kDa cut-off to yield only digitiferin. Flow-through and elution fractions were collected and analyzed by Tris-Tricine SDS-PAGE and gel staining with Coomassie Brilliant Blue R-250 (NACALAI TESQUE, Kyoto, Japan). The protein concentration was measured using the Bradford protein assay (Bio-Rad Laboratories, California, USA), with bovine serum albumin as a standard.

**Determination of antimicrobial activity against *E. coli***
*E. coli* cells pre-cultured in LB medium ($OD_{600}$ = 0.1) were collected and inoculated into PB (Poor broth: 1% [w/v] tryptone) at 30℃ for 21h. *E. coli* grown to exponential phase were collected at 1.0 × $10^3$ CFUs/mL and 100 µL were added to a glass-bottomed dish (IWAKI, Tokyo, Japan). Purified digitiferin was adjusted to a final concentration of 10 µM using 20 mM HEPES buffer (pH 7.0) and incubated with the bacterial suspension at 30℃ for 18 h. Trypan blue (Fujifilm Wako Pure Chemical Corporation, Osaka, Japan) was added to determine whether cells were alive or dead. Cells were observed using an all-in-one fluorescence microscope (BZ-X810; Keyence, Osaka, Japan).

**Evaluation of the antimicrobial spectrum of digitiferin**
Using a liquid growth inhibition assay, antibacterial activity was assayed against Gram-negative bacteria (*E. coli* [NBRC 102203$^T$], *P. aeruginosa* [NBRC 12689$^T$], and *S. marcescens* [ATCC BAA-632]), Gram-positive bacteria (*B. subtilis* [NBRC 13719$^T$] and *S. aureus* [NBRC 100910$^T$]) and coral-pathogenic bacteria (*V. coralliilyticus* P1 [LMG23696], *V. coralliilyticus* YB1 [ATCC BAA-450], *V. shiloi* AK1 [ATCC BAA-91] and *S. marcescens* [ATCC BAA-632]). Gram-negative and Gram-positive bacteria pre-cultured in LB medium ($OD_{600}$ = 0.1) were collected, and inoculated into PB. The pre-culture medium was MB for coral-pathogenic bacteria and they were used to inoculate PB supplemented with NaCl (PB-NaCl; 1.5% [w/v] NaCl). Bacteria cultured to logarithmic growth phase were adjusted to 1.0 × $10^4$ CFUs/mL and 100 µL were added to wells of 96-well plates (Thermo Fisher Scientific). Purified digitiferin was diluted in a series of 2-fold dilutions from 10 to 1.25 µM with 20 mM HEPES buffer (pH 7.0), and aliquots (20 µL) were added to the bacterial suspension. The plate was incubated at 30°C for 18 h, and the $OD_{600}$ was measured using a microplate reader (SpectraMax iD3; Molecular Devices LLC, San Jose, CA, USA). After incubation, MICs were recorded as the smallest dilution that inhibited bacterial growth (measured at $OD_{600}$). MBCs (≧99.9% killing) were determined by plating contents of the first three wells with no visible bacterial growth onto LB agar plates for Gram-negative and Gram-positive bacteria and MA plate for coral-pathogenic bacteria. The lowest concentration of digitiferin that prevented colony formation was recorded as the MBC. MACs were the lowest concentration of digitiferin-treated bacteria that formed aggregates. The suspension of the medium was evaluated visually.

**3D structural model of digitiferin**
The 3D structure of digitiferin (residues 20-100) was predicted using ColabFold (5). The structural model was viewed with PyMOL software. The structure of digitiferin was analyzed using APBS to

calculate macromolecular electrostatics (6). Using the DALI server, we then searched for structural similarities between digitiferin and known proteins (http://ekhidna2.biocenter.helsinki.fi/dali/).

**SEM observations**

Digitiferin-treated bacteria were observed using SEM. *B. subtilis* was pre-cultured overnight in LB medium and inoculated with PB. *B. subtilis* grown to exponential phase were adjusted to 1.0 × $10^4$ CFUs/mL and incubated with digitiferin at final concentrations of 5 μM at 30°C for 1 h to clarify the effects of digitiferin on bacteria. Bacteria were incubated with 20 mM HEPES buffer (pH 7.0) as a negative control. Subsequently, fixing solution (1% [v/v] glutaraldehyde; 1% [w/v] NaCl; 0.1 M cacodylate buffer) was added to the bacterial suspension and incubated at 4℃ overnight. The treated cell suspension was loaded onto a nano-percolator filter (JEOL Ltd., Tokyo, Japan) and aspirated with a 5-mL syringe attached to the filter holder. Bacterial samples on the filter were washed twice with 0.1 M cacodylate buffer and dehydrated using an ethanol series and tert-butyl alcohol. After substituting t-butyl alcohol at -4°C, samples were dried using a freeze-drier (JFD-300; JEOL Ltd.) overnight. Platinum-palladium coating of dried samples was performed using an ion sputter (E-1030; Hitachi, Tokyo, Japan) and samples were observed by a Scanning Electron Microscope (S-4800; Hitachi).

**Determining the specificity of anti-digitiferin**

For histological characterization of cells expressing digitiferin, a polyclonal antibody against digitiferin (anti-digitiferin antibody) was developed using 7.0 mg of purified digitiferin as an antigen (Biologica, Nagoya, Japan). To determine the specificity of anti-digitiferin, Western blotting was performed on a recombinantly expressed *E. coli* crude sample. As a result, a single band of the target size (25.5 kDa) was detected (data not shown).

The specificity of this antibody to the purified antigenic peptide by dot blot analysis was examined. Purified digitiferin was dotted on a nitrocellulose membrane (Fujifilm Wako Pure Chemical Corporation) in the amounts of 10, 1, 0.1 and 0 ng. After washing with TBS containing 0.1% Tween 20 (TBST), membranes were incubated in anti-digitiferin antibody (diluted from 1:1000 to 1:32,000 with TBST containing 2% skim milk). After washing with TBST, alkaline phosphatase-conjugated goat anti-guinea pig IgG antibody (abcam, Cambridge, UK; diluted 1:4,000 in TBST containing 2% skim milk) was used for the secondary anti-body reaction. Immunoreactivity was visualized using the NBT/BCIP liquid substrate system (Sigma-Aldrich, Missouri, USA).

**Histological analysis**

Corals were collected and cultured as previously described (7). Permits for coral collection were obtained from the Okinawa Prefectural Government (Permit no. 4-33) for research use. Coral fragments were fixed for 16 h at room temperature with a fixing solution containing Zinc Formalin Fixative (pH 6.25; Polysciences Inc., Warrington, PA), 0.2 % glutaraldehyde (Fujifilm Wako Pure Chemical Corporation), and 35 g/L Daigo’s artificial seawater (Nihon Pharmaceutical, Tokyo, Japan). The fixing solution was improved on a previous study (8). Samples were decalcified with 0.5 M EDTA (pH 8.0; Nippon Gene Co. Ltd., Toyama, Japan) for 5 days and dehydrated sequentially through 70, 80, 90, 95, 100 % ethanol (100% ethanol was changed twice). After dehydration, samples were replaced in xylene twice and embedded in paraffin. Tissue sections were cut at a thickness of 4 μm each. In this study, > 3 colonies of *A. digitifera* were analyzed.

Immunohistochemistry was performed according to a previous study (9). Sections were dewaxed in xylene twice and put in 100, 95, 90, 80, 70 % ethanol (100% ethanol was changed twice). After washing with PBS containing 0.1% Tween 20 (PBST), For the primary antibody reaction, sections were incubated with anti-digitiferin antibody diluted 1:6,000 in PBST with 5 % skim milk for 16 h at 4°C. As negative controls, a guinea pig serum IgG from a non-immunized animal (Fujifilm Wako Pure Chemical Corporation) and anti-digitiferin antibody pre-adsorbed with 100 µg/mL of the peptide antigen were used. After washing with PBST, sections were incubated with the secondary antibody, a biotinylated goat anti-guinea pig IgG antibody (Vector Laboratories, Burlingame, CA; diluted 1:4,000). After washing with PBST, sections were then incubated in standard avidin-biotinylate-peroxidase complex solution (ABC kit, Vector Laboratories) for 30 min. Immunoreactivity was visualized using 3,3'-diaminobenzidine (Sigma-Aldrich). Cell nuclei were stained with haematoxylin (MUTO PURE CHEMICALS CO, Tokyo, Japan). Sections were observed and photographed with an all-in-one fluorescence microscope (BZ-X810; Keyence).

**Salt sensitivity of digitiferin**

To analyze the salt sensitivity of digitiferin, a liquid growth inhibition assay was performed using *B. subtilis.* As described above, *B. subtilis* pre-cultured in LB medium overnight was inoculated onto normal PB or supplemented with NaCl (final concentrations of 0.1, 0.25, 0.5, 1.0% [w/v]). *B. subtilis* was adjusted to $1.0 \times 10^4$ CFUs/mL and 100 µL were added to 96-well plates. Purified digitiferin was diluted in a series of 2-fold dilutions from 10 to 1.25 µM with 20 mM HEPES buffer (pH 7.0). Aliquots (20 µL) were added to the bacterial suspension. The plate was incubated at 30°C for 18 h, and the $OD_{600}$ was measured.

**Sodium ion measurement of coral mucus**

Colonies of *A. digitifera* were exposed on the surface at low tide, and their secreted mucus collected by syringe aspiration. Sodium ion concentrations of coral mucus and seawater were measured using a measuring device (LAQUAtwin Na-11; Horiba, Kyoto, Japan).

**Antibacterial assay for coral-pathogenic bacteria with mannitol medium**

As when evaluating the antimicrobial spectrum, coral-pathogenic bacteria were pre-cultured in MB and maintained at logarithmic growth phase in PB-NaCl. Bacteria were adjusted to $1.0 \times 10^4$ CFUs/mL with PB-mannitol (18.5% [w/v] mannitol). Purified digitiferin was diluted in a series of 2-fold dilutions with 20 mM HEPES buffer (pH 7.0) and aliquots (20 µL) were added to the bacterial suspension (100 µL). The plate was incubated at 30°C for 3 h. All wells were plated onto MA plates, and antimicrobial activity was evaluated by counting CFUs after incubation at 30°C for 24 h.

**Statistical analysis**

All results are given as means ± standard error of the mean (n = 3). Statistical analysis for antibacterial assays were conducted using t-test in EZR software (Jichi Medical University Saitama Medical Center, Saitama, Japan; http://www.jichi.ac.jp/saitama-sct/SaitamaHP.files/statmedEN.html) (10), which is a graphical user interface for R (R Foundation for Statistical Computing, Vienna, Austria, v2.13.0). *P* values less than 0.05 were considered statistically significant.

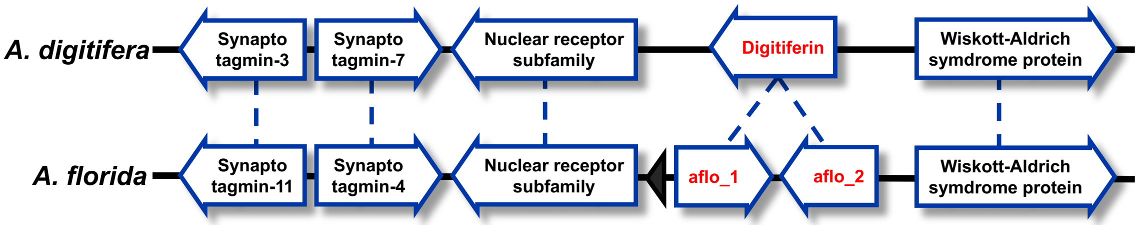


Fig. S1 Conserved syntenic region of genes around digitiferin in *Acropora digitifera* (scaffold 0027) and *Acropora florida* (scaffold 0063). The gene encoding digitiferin was duplicated in *A. florida* (digitiferin: adig_s0027.g47 from *A. digitifera*; aflo_1: aflo_s0063.g39 from *A. florida;* and aflo_2: aflo_s0063.g40 from *A. florida).* Arrows indicate the transcription direction. Possible gene names are also shown in boxes. Syntenic regions of *A. digitifera* and *A. florida* are conserved with two genes encoding synaptotagmin, which promotes synaptic vesicle exocytosis.

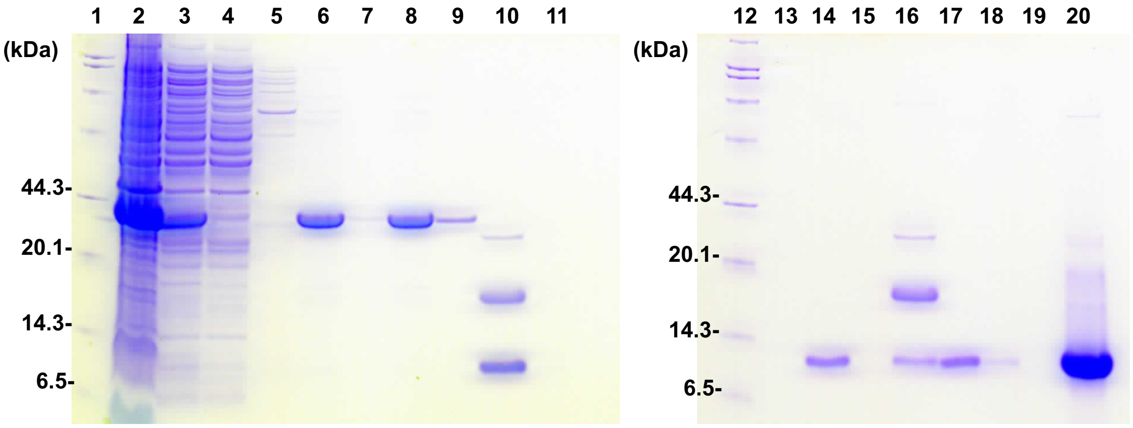


Fig. S2 Coomassie brilliant blue-stained analytical SDS-PAGE showing IPTG induction of recombinant digitiferin expressed in *E. coli* and purification by Ni-affinity chromatography. Lanes 1 and 12, low-molecular weight markers; lane 2, insoluble fraction; lane 3, soluble fraction (the molecular weight of recombinant digitiferin [25 kDa]), lane 4 to 6, flow-through eluted with buffer included 5 mM, 20 mM, or 200 mM imidazole, lane 7, centrifuged precipitation of lane 6; lane 8, dialyzed of lane 6; lane 9, centrifuged precipitation of lane 8; lane 10, adding to HRV 3C protease (the molecular weight of HRV 3C protease [22 kDa], thioredoxin-6×His [15.9 kDa] and digitiferin [9.6 kDa]), lane 11, centrifuged precipitate of lane 10; lanes 13, 14 and 16, flow-through eluted with buffer included 5 mM, 20 mM or 200 mM imidazole; lane 15, centrifuged precipitate of lane 14; lane 17, dialyzed of lane 14; lane 18, centrifuged precipitate of lane 17; lane 19, centrifuged precipitate of lane 20; lane 20, concentrated of lane 17 (the molecular weight of digitiferin [9.6 kDa]).

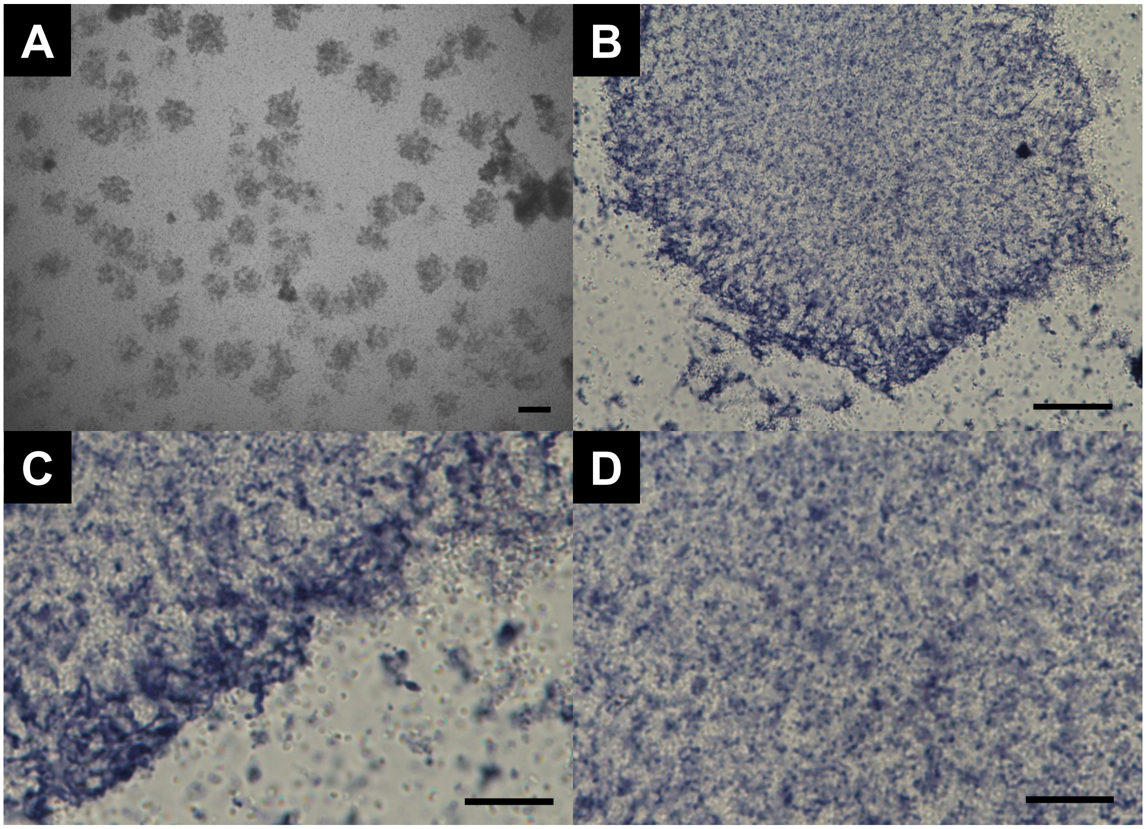


Fig. S3 Bacterial aggregates of *E. coli* treated with digitiferin. (A) *E. coli* formed aggregates at 10 μM digitiferin. (Scale bar, 200 μm) (B) Trypan-blue staining of aggregation. Only dead cells were stained blue. (Scale bar, 20 μm) (C) Edges of the aggregate. (Scale bar, 10 μm) (D) Aggregate centers contained both live and dead cells. (Scale bar, 10 μm)

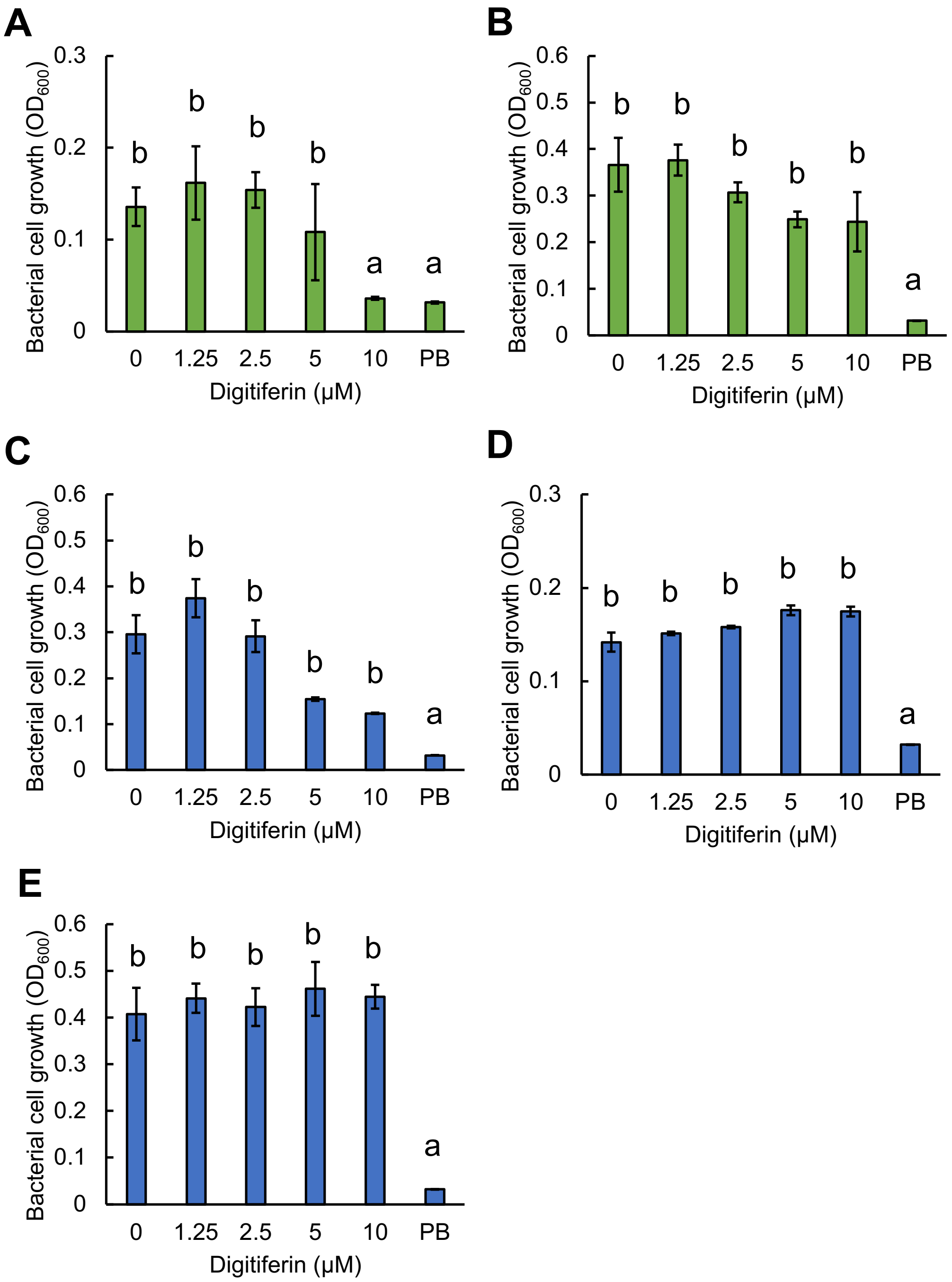


Fig. S4 Growth of digitiferin-treated Gram-positive and Gram-negative bacteria. Antimicrobial activity against Gram-positive and negative bacteria was evaluated using a liquid growth inhibition assay. Bacterial growth was measured using a microplate reader ($OD_{600}$) to determine MICs. The digitiferin concentration was varied from 0 to 10 μM. PB: poor broth without bacteria. All data are presented as means ± standard error of the mean, $n = 3$. Statistical analysis between each sample and the negative control (PB) was conducted using t-test. Significant differences are represented by different letters. (A) *B. subtilis* (B) *S. aureus* (C) *E. coli* (D) *P. aeruginosa* (E) *S. marcescens*

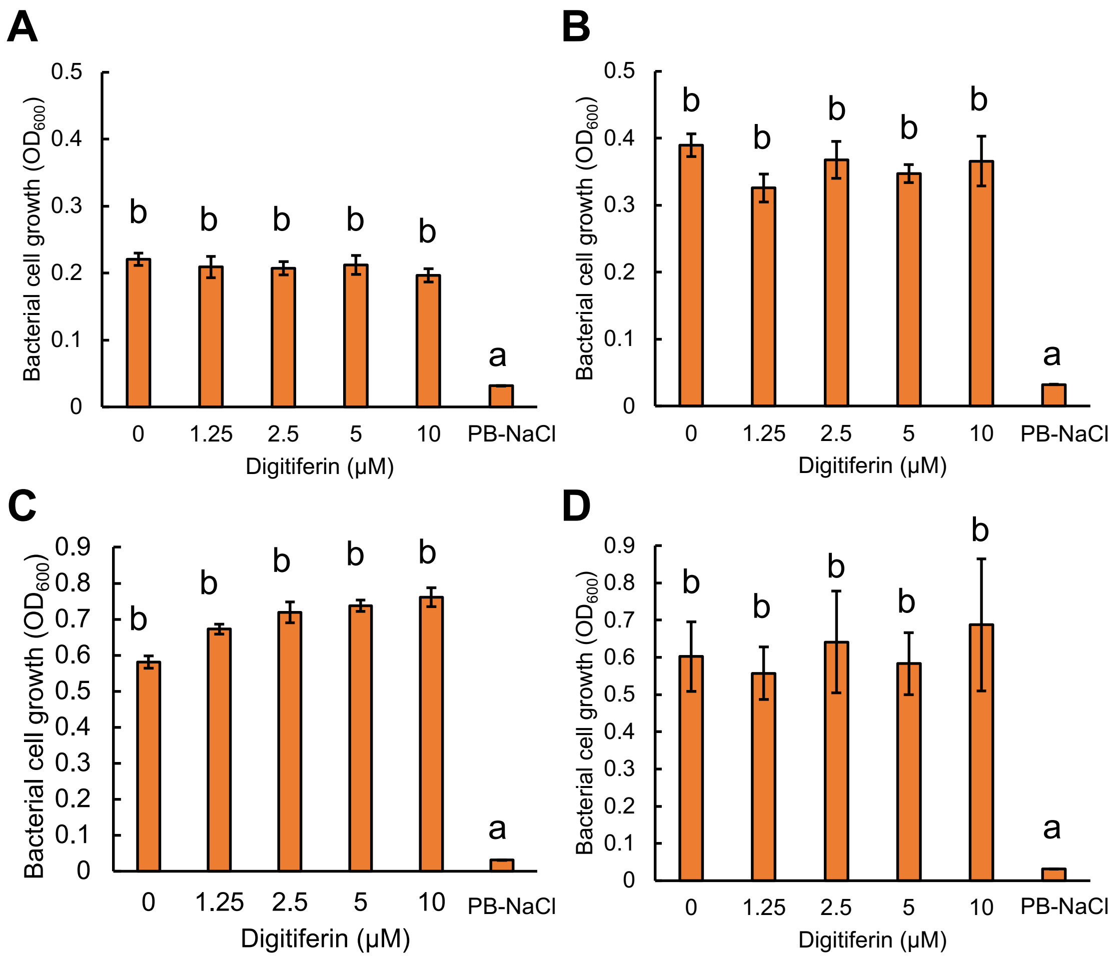


Fig. S5 Growth of digitiferin-treated coral-pathogenic bacteria. A liquid growth inhibition assay evaluated antimicrobial activity against coral-pathogenic bacteria. Bacterial growth was measured using a microplate reader ($OD_{600}$) to determine MICs. The digitiferin concentration was varied from 0 to 10 μM. PB-NaCl, poor broth supplemented with 1.5% (w/v) NaCl without bacteira. All data are presented as means ± standard error of the mean, $n$ = 3. Statistical analysis between each sample and the negative control (PB-NaCl) was conducted using t-test. Significant differences are represented by different letters. (A) *V. coralliilyticus* P1 (B) *V. coralliilyticus* YB1, (C) *V. shiloi* AK1, and (D) *S. marcescens*

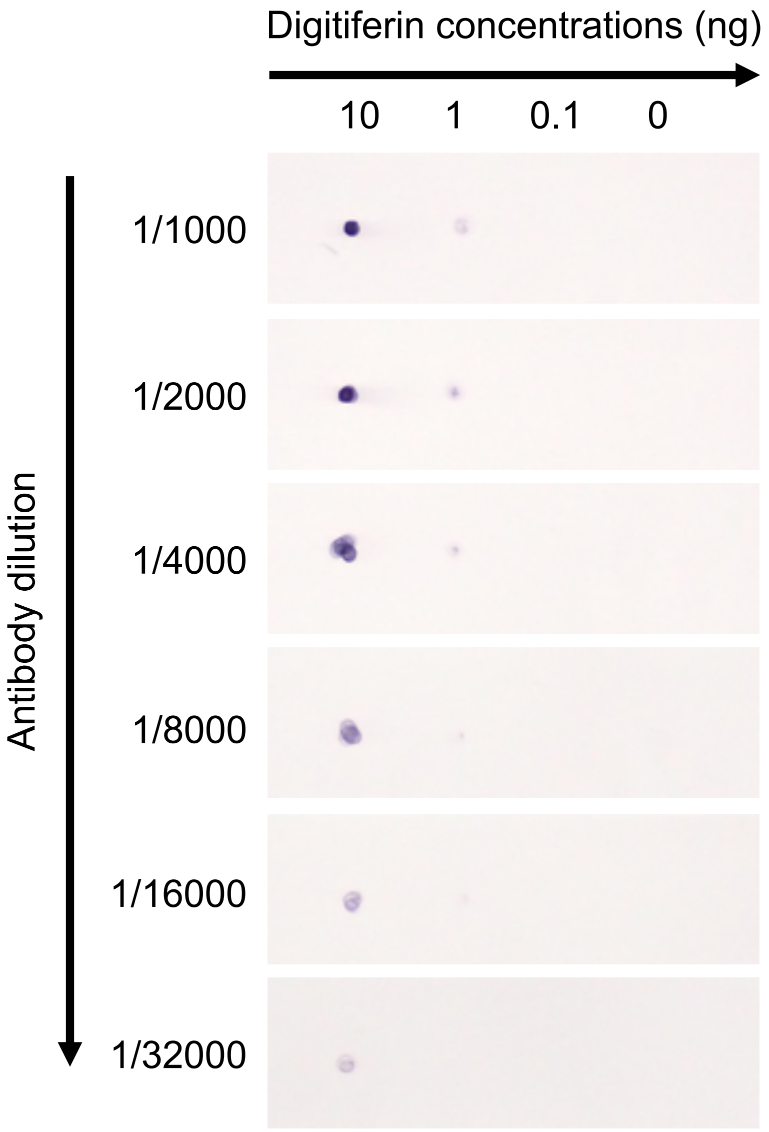


Fig. S6 Evaluation of the specificity of anti-digitiferin antibody by dot blot analysis. The peptide antigen used for antibody production was dotted on the nitrocellulose membrane in amounts of 10, 1, 0.1, and 0 ng. Antibody dilution factors ranged from 1:1000, 1:2000, 1:4000, 1:8000, 1:16000, and 1:64000.

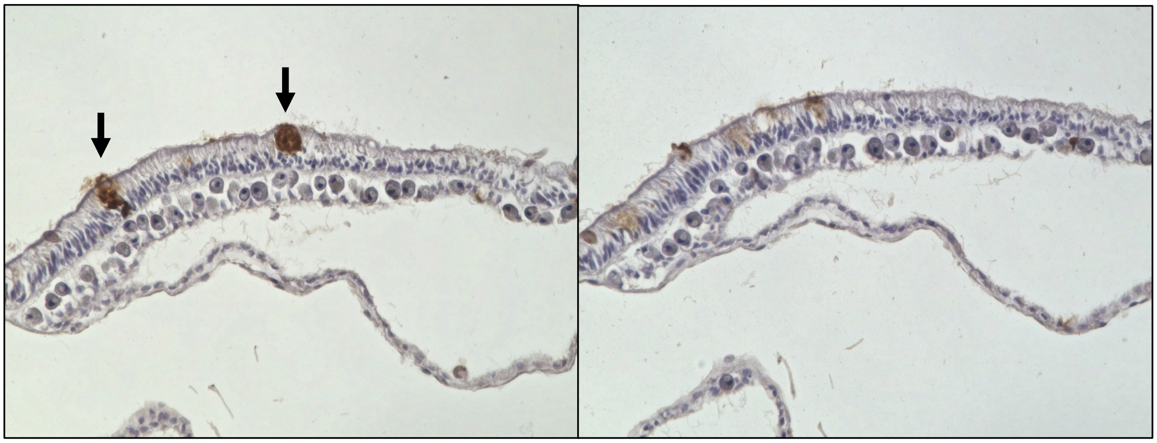

Fig. S7 Immunohistochemical analysis with anti-digitiferin antibody and anti-digitiferin antibody pre-adsorbed with the antigenic peptide (pre-adsorbed). (A) Immunohistochemistry with anti-digitiferin antibody. Arrows indicate localization of digitiferin. (B) Immunohistochemistry with pre-adsorbed antibody, as negative control. The anti-digitiferin antibody was pre-adsorbed with 100 μg/mL of the antigenic peptide and used in place of the primary antibody. Pre-adsorption resulted in substantially decreased signals. Sequential sections were used for the analysis.

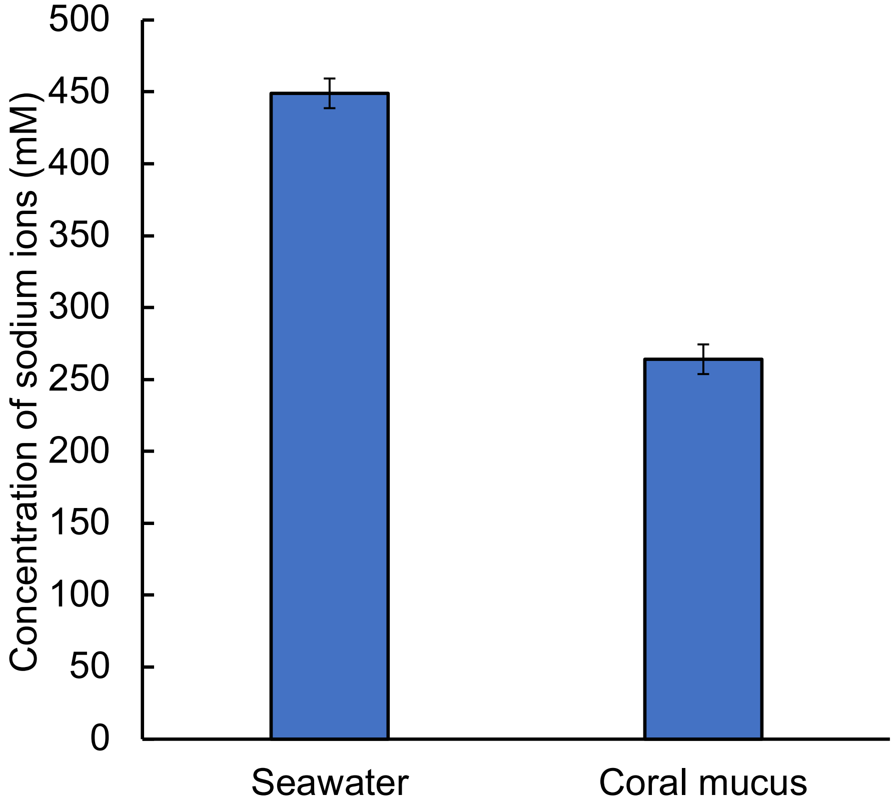


Fig. S8 Concentration of sodium ions in coral mucus and surrounding seawater. The sodium ion concentration of coral mucus was 264 mM. In comparison, seawater at Sesoko Island, the sampling site of *A. digitifera*, was 449 mM. All data are presented as means ± standard error of the mean, $n$ = 3.

**Table S1 Primer sequences used for PCR.**

| Name | Orientation | Sequence (5'-3') |
|---|---|---|
| digitiferin | Forward | GCAAACCAGGCAAAGTAAATCAGTT |
| | Reverse | TGATTGTTACGGTATTTGCACTTCA |
| M13 | Forward | CAGGAAACAGCTATGAC |
| | Reverse | GTTTTCCCAGTCACGAC |
